# Additive and non-additive epigenetic signatures of hybridisation between fish species with different mating systems

**DOI:** 10.1101/2020.07.01.182022

**Authors:** Waldir M. Berbel-Filho, Andrey Tatarenkov, George Pacheco, Helder M. V. Espírito-Santo, Mateus G. Lira, Carlos Garcia de Leaniz, John C. Avise, Sergio M. Q. Lima, Carlos M. Rodríguez-López, Sofia Consuegra

## Abstract

Hybridisation is a major source of evolutionary innovation. However, several prezygotic and postzygotic factors influence its likelihood and evolutionary outcomes. Differences in mating systems can have a major effect on the extent and direction of hybridisation and introgression. In plants, epigenetic mechanisms help to stabilize hybrid genomes and contribute to reproductive isolation, but the relationship between genetic and epigenetic changes in animal hybrids is unclear. We analysed the extent of a unique case of natural hybridisation between two genetically distant mangrove killifish species with different mating systems, *Kryptolebias hermaphroditus* (self-fertilising) and *K. ocellatus* (outcrossing), and the methylation patterns of their hybrids. Hybridisation rate between the species ranged between 14% and 26%. Although co-existing parental species displayed highly distinct genetic (microsatellites and SNPs) and methylation patterns (37,000 differentially methylated cytosines), our results indicate that F1 hybrids are viable and able to backcross with parental species. Hybrids had predominantly intermediate methylation patterns (88.5% of the sites) suggesting additive effects, as expected from hybridisation between genetically distant species. Differentially methylated cytosines between hybrids and both parental species (5,800) suggest that introgressive hybridisation may play a role in generating novel genetic and epigenetic variation which could lead to species diversification. We also found a small percentage of non-additive epigenetic effects which might act as an evolutionary bet-hedging strategy and increase fitness under environmental change.

## Introduction

Hybridisation is a major source of evolutionary innovation, with important implications for phenotypic diversification, adaptation and ultimately speciation (1, 2). Hybridisation can result in heterosis or hybrid vigour, with hybrids displaying higher performance than the parents (3, 4), but also in hybrid incompatibility resulting in loss of fitness, increased mortality, and reproductive isolation (5, 6). Hybridisation is particularly high in plants (40%) (7, 8), insects and birds (10%) (9). In fish like minnows (genus *Gila*) and killifish (*Fundulus grandis*), introgressive hybridisation has played a central role in their diversification and/or adaptation, by introducing additional genetic variation for selection and drift to act upon (10, 11). Yet, the prezygotic factors influencing the likelihood of interspecific hybridisation, and the postzygotic molecular mechanisms underlying its outcomes, remain poorly understood (12, 13).

The strength of prezygotic barriers is a major factor keeping reproductive isolation (14). Genetic, spatial, temporal, and behavioural differences can help to maintain sympatric species reproductively isolated (15). In addition, differences in mating systems have particularly strong effects on the likelihood, extent and direction of hybridisation and introgression. In monkeyflowers (genus *Mimulus*), for example, differences in the mating system (predominantly selfing vs obligate outcrossing) have an overriding effect over other prezygotic and postzygotic mechanisms, leading to complete reproductive isolation (16). Unlike plants, most animals are dioecious (17, 18) and relatively little is known about how mating systems could act as prezygotic barriers for hybridisation in animals when compared to plants (14, 16, 19).

Hybrid incompatibility (HI), encompassing hybrid unviability, sterility and reduced fitness, is one of the strongest postzygotic mechanisms acting as isolating barriers for interspecific hybridisation (20). HI was initially thought to be caused solely by interactions between incompatible parental genetic alleles (21, 22), but can also result from changes in regulatory elements (23), transposable elements activity (24) (Ungerer et al. 2006), chromosomal rearrangements (25), cytosine methylation (13, 26) and changes in gene expression patterns (27, 28). For instance, a single epigenetically inactivated gene (HISN6B) is responsible for hybrid incompatibility between strains of *Arabidopsis thaliana* (5). Interactions between parental genomes can result in additive or non-additive gene expression patterns (29). Thus, hybrids between farmed and wild Atlantic Salmon or between recently-diverged pupfish species have shown mostly non-additive patterns of gene expression (e.g. over or under-dominance) relatively to the parental species (27, 30), while hybrids of house mouse subspecies and *Drosophila* species display predominately additive effects (31, 32), suggesting that additivity in gene expression can be expected in more divergent taxa (31). The epigenetic effects of hybridisation are much less known, particularly in animals. Epigenetic mechanisms, particularly DNA methylation, play a central role in the initial stabilization of the genome of allopolyploid plant hybrids and their evolutionary success, by for example gene silencing and dosage compensation (33). However, it is unclear to what extent the role of epigenetic modifications in the restructuring of the hybrid genomes is determined by the underlying genetic changes (34). In plant hybrids, both additive (35) and non-additive (36) effects can modify DNA methylation patterns, the direction of which depends on the initial degree of divergence between parental lineages (37). For example, hybrids between inbred lines of *Arabidopsis* display non-additive changes in DNA methylation more frequently when parental methylation levels are different, but not when they are similar (38).Thus, epigenetic changes during hybridisation are related to changes in gene regulation and potentially to reproductive isolation which could lead to speciation, but the extent to which these changes are dependent from the genetic background and the distance between the parental species is unclear, particularly in animals (34).

Here, we analysed the extent and direction of a unique case of natural hybridisation, recently identified, between two genetically distant mangrove killifish species with different mating systems, *Kryptolebias hermaphroditus* (self-fertilising) and *K. ocellatus* (outcrossing) (39) (Supplementary material). We also analysed the epigenetic (methylation) patterns of the hybrids and compared them to the parental species’, to determine the extent to which they are determined by the genomic background.

## Results

### Hybridisation between K. hermaphroditus and K. ocellatus

Both SNPs (Fig. 1) and microsatellite genotypes (Fig. S3) confirmed the high rates of *K. hermaphroditus* and *K. ocellatus* hybridisation. Microsatellites identified six hybrids from Guanabara Bay (Fundão, FUN: FUN 08, 11, 13, 41, 43, 47; 26% of the individuals sampled) and five from Sepetiba Bay (Guaratiba, GUA: GUA 09, 17, 20, 24, 62; 14% of the individuals sampled in 2017; Fig. S3) with unique alleles at nine loci, not present in any other *K. ocellatus* but fixed in *K. hermaphroditus* from FUN and GUA populations. STRUCTURE analysis also supported the identification of these hybrids (Fig S3). At K = 2 (most likely partition according to Evanno’s ΔK) all *K. hermaphroditus* individuals were assigned with nearly 100% probability to one cluster, and almost all *K. ocellatus* to another cluster, with exception of the subset of the divergent FUN and GUA fish, with admixed genetic backgrounds of both species. At K = 3 the southernmost *K. ocellatus* individuals (SFR and FLO) formed their own genetic cluster (Fig. S3), reflecting the deep genetic structuring previously found between southeast and south populations on this species.

**Figure 1.**
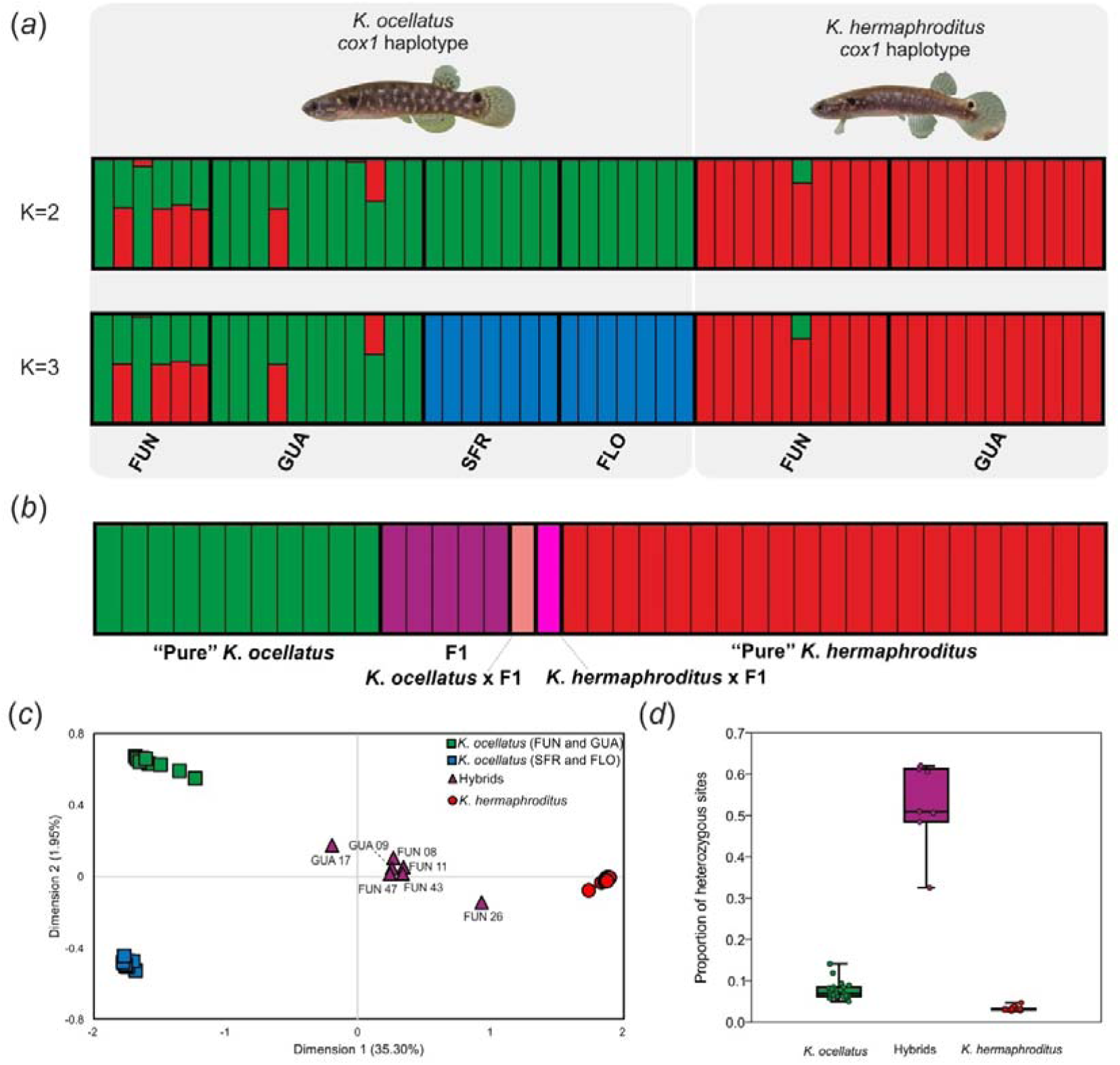
Genetic clustering, diversity and hybrid classification across *K. ocellatus* and *K. hermaphroditus* individuals (*a*) Admixture plots for K= 2 and K=3 based on 5,477 SNPs extracted from 53 *Kryptolebias spp.* individuals generated using ngsAdmix v.3 .2. Each individual is represented by a bar, and each colour represents a genetic cluster. (*b*) NEWHYBRIDS individual classification using a subset of 200 SNPs with low-levels of linkage-disequilibrium. (*c*) Multidimensional Scaling (MDS) based on the genetic distances from 5,477 SNPs. Hybrid individuals (see results) are highlighted with their respective labels. (*d*) Proportion of heterozygous sites between *K. ocellatus, K. hermaphroditus* and the hybrid individuals.

STRUCTURE results using K = 2 indicated that about half of the genetic ancestry of most of the hybrid fish from FUN (FUN 08, 11, 13, 43, 47) belonged to *K. hermaphroditus* (average q-value for *K. hermaphroditus* cluster = 0.58, sd ± 0.01), with exception of FUN41, with only 18% of *K. hermaphroditus* ancestry. In GUA, the ancestry of two individuals (GUA 09 and 62) was about 50% from *K. hermaphroditus* (average q-value = 0.51, sd ± 0.02), while the other three potential hybrids (GUA 17, 20, 24), displayed approximately 35% of *K. hermaphroditus* ancestry (average q-value = 0.35, sd ± 0.02). This was supported by the results of NEWHYBRIDS, with seven potential hybrids (FUN 08, 11, 13, 47, 48; GUA 09 and 62) being classified as F1 from outcrossing between *K. ocellatus* and *K. hermaphroditus* (average probability a posteriori = 0.97, sd ± 0.05), and four (FUN 41; GUA 17, 20, 24) as backcrosses between *K. ocellatus* and a F1 hybrid (average probability a posteriori = 0.99, sd ± 0.003). No individuals were identified as F2 or as backcrosses involving *K. hermaphroditus* (Fig. S3). All hybrids had a *cox1* haplotype typical of *K. ocellatus*, suggesting that this species would act as the female part in crosses with *K. hermaphroditus* males (Fig. S3).

Admixture results based on 5,477 SNPs were largely consistent with the genetic structure and hybridisation evidence provided by microsatellite data, with six fish displaying strong evidence of admixture. At K = 2, four individuals from FUN with *K. ocellatus cox1* haplotypes (FUN 08, 11, 43, 47) and two from Guaratiba (GUA 09, 17) showed a mixed ancestry between *K. ocellatus* and *K. hermaphroditus* (Fig. 1a). Five additional individuals identified as hybrids by microsatellite data could not be classified, two from FUN (FUN 13, 41) which failed to produce enough reads for the GBS library (cut-off ≥ 500k reads) and three from GUA (GUA 20, 24, 62) which were not included in the library (Table S2). One individual from Fundão (FUN 26) showed genetic admixture only with SNPs. As for the microsatellites at K = 3, *K. ocellatus* from southeastern and southern populations were split in separate clusters (Fig 1a). Results from NEWHYBRIDS using a subset of 200 SNPs with high F_ST_ and low LD classified five individuals (FUN 08, 11, 43, 47; GUA 09) as F1 (probability *a posteriori* equals 1 in all individuals) and one individual (GUA 17) as a backcross between *K. ocellatus* and a F1 hybrid (probability *a posteriori* equals 1). These results agreed with the microsatellites, with the exception of FUN 26, classified as *K. hermaphroditus* with microsatellites but as a backcross between *K. hermaphroditus* and a F1 hybrid (posterior probability=1) with SNPs. No individual was classified as F2 (Fig. 1b). Hybrids (FUN 08, 11, 26, 43, 47; GUA 09, 17) had a mean proportion of heterozygous sites of 0.53, approximately ten-fold the values of *K. ocellatus* (0.07) and *K. hermaphroditus* (0.03) (Fig. 1d).

### Cytosine methylation patterns in hybrids and parental species

The msGBS library only including individuals from FUN and GUA yielded in average 6,422,972 reads per individual, with 85.21% reads uniquely mapping to *K. marmoratus* reference genome (ASM164957v1) (Table S2) corresponding to 830,905 loci.

In total, 37,664 significant differentially methylated cytosines (DMCs) were found between *K. ocellatus* and *K. hermaphroditus* (false discovery rate < 0.01). *K. hermaphroditus* showed higher number of DMCs than *K. ocellatus* relative to the hybrids (hybrids vs *K. hermaphroditus*: 13,905 DMCs; hybrids vs *K. ocellatus*: 10,620 DMCs) (Fig. 2a), with 50.45% of the DMCs hypermethylated in *K. hermaphroditus* in relation to the hybrids (and vice-versa for hypomethylated DMCs) compared to 64.23% in *K. ocellatus* (Fig. 2a). MDS analysis using DMCs between parental species positioned the hybrids between two clusters representing the parental species (Fig. 2d). These results were also supported by the MDS using all reads normalised by library size (830,905 sites) (Fig. S5). In both cases, individuals identified as backcrosses occupied eigen spaces closer to the parental species than to F1 hybrids.

**Figure 2.**
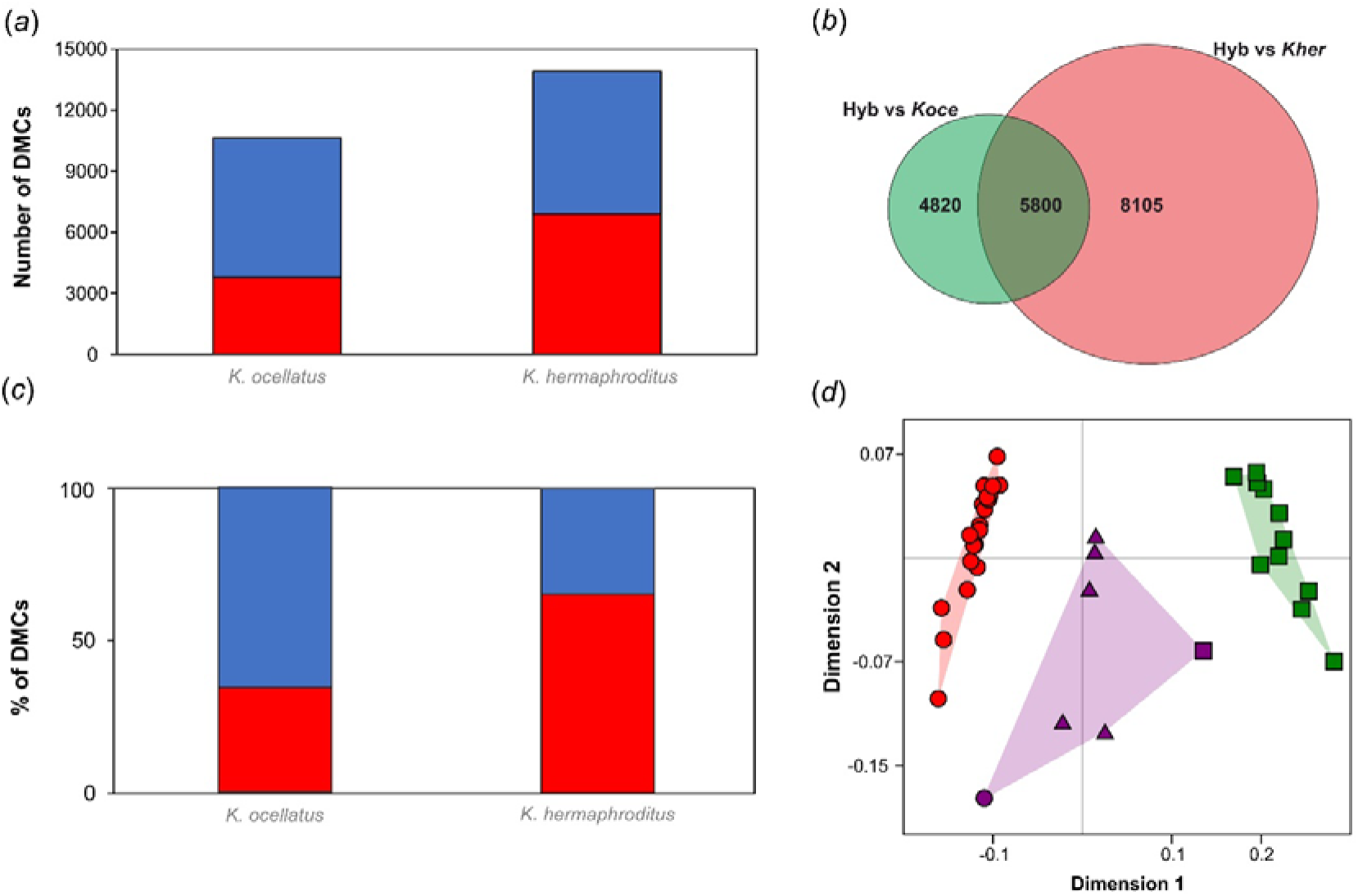
Cytosine methylation comparisons between parental species and hybrids. *(a*) Number of differentially methylated cytosines (DMCs) in *K. ocellatus* and *K. hermaphroditus* compared to their hybrids. Hypomethylated (logFC value > 1) and hypermethylated (logFC value < 1). DMCs in comparison to hybrids are shown in blue and red, respectively. (*b*) Overlap in the number of DMCs of *K. ocellatus* and *K. hermaphroditus* in comparison with their hybrids. (*c*) Percentage of DMCs either hypomethylated (blue) or hypermethylated (red) of the 5,800 DMCs common to the comparisons between hybrids vs parental species. (*d*) Multidimensional scaling analyses of the normalised counts for the 5,800 DMCs common to the comparisons between hybrids vs parental species. Squares represent *K. ocellatus*, circles represent *K. hermaphroditus*, and triangle represent hybrids. Backcrosses are represented by purple shapes according to their respective parental species.

Hybrids shared 5,800 common DMCs with both parental species (Fig. 2b), of which 34.28% and 65.72% were hyper and hypo methylated, respectively, in *K. ocellatus* in relation to the hybrids (65.19% and 34.81% in *K. hermaphroditus*; Figs.2c and S6). The MDS analysis showed two parental clusters and an intermediate cluster containing the hybrids (both F1 and backcross individuals). Backcrossed individuals, GUA 17 and FUN 26, occupied eigen spaces closer to the parental species than F1 hybrids (Fig. 2d).

All hybrids displayed intermediate levels of DNA methylation relatively to the parental species in the hierarchical clustering analysis, with exception of the backcrossed individuals, GUA 17 and FUN 26, which clustered with their respective parental species, *K. ocellatus* and *K. hermaphroditus* (Fig. S7). Of the 37,664 DMCs between *K. ocellatus* and *K. hermaphroditus*, 33,329 (88.5%) had intermediate normalised read counts in the hybrids compared to the parental species. The remaining 4,435 (11.5%) DMCs had either higher (77.3%) or lower (22.7%) number of normalised reads in hybrids when compared to parental species. Generally, the same pattern was observed in the analysis of the 5,800 DMCs common in the comparisons between hybrids vs parental species, with 98.2% showing intermediate number of reads.in hybrids, while 1.8% were either over or under dominant on hybrids (65 and 34 DMCs, respectively).

### Gene ontology analysis

Of the 5,800 DMCs shared between hybrids and parental species, 254 (4.38%) were within 2kb upstream gene bodies, representing putative promoters, 3,435 (59.23%) were overlapping gene bodies, while 831 (14.33%) represented potential intergenic regions. Of these, 1,280 (22.06%) were within unannotated regions and the rest affected putative promoters and/or gene bodies of 2,786 unique genes, 1,322 of them mapping to orthologs in the zebrafish genome. The gene ontology enrichment analysis identified 217 significantly overrepresented ontologies influencing a wide range of biological processes, from cellular regulation, to neurogenesis and organ development (Table S3). The MDS analysis separating DMCs by genomic context revealed a clear clustering between parental species across all genomic contexts (e. g. promoters, gene bodies, intergenic and unannotated regions), with the hybrids forming an intermediate cluster differentiated from the parental species (Fig. S8).

## Discussion

Hybridisation is expected to be rare between species with different mating systems because different modes of reproduction should act as strong prezygotic barriers (4). Yet, evidence for this in animals is lacking. Our results revealed widespread hybridisation and introgression between two mangrove killifish species with different mating systems, *K. hermaphroditus* (predominately selfing) and *K. ocellatus* (obligately outcrossing) at unusually high rates (up to 26%). Hybrids between these species displayed high heterozygosity and predominantly Mendelian-like epigenetic inheritance patterns, with intermediate levels of cytosine methylation relative to the parental species, indicative of additive effects, reflecting a strong influence of the genetic background on cytosine methylation (40, 41).

Hybridisation has previously been observed in *Kryptolebias* (Tatarenkov et al. 2018; Tatarenkov et al. 2020), albeit in more closely related species (3% K2P distance at *cox1* between compared to 11% in this case) with similar mating system, and could represent an important source of novel genetic and epigenetic variation, particularly for the selfing species. In the selfing and predominantly inbred mangrove killifish species *K. marmoratus* and *K. hermaphroditus*, genetic diversity is also known to increase by occasional male-mediated outcrossing (40, 42). As *K. marmoratus* males prefer to associate with genetically dissimilar hermaphrodites (43) and increased heterozygosity is related to lower parasite loads (40, 42), it is likely that mechanisms that increase genetic diversity are important for the fitness and survival of the species. Genetic evidence suggests that hybridisation may be relatively recent within this system resulting from the recent colonisation of *K. hermaphroditus* of an area already inhabited by *K*. ocellatus (39, 44); this could have helped the range expansion *K. hermaphroditus*, as it has been observed for other invasion processes (3).

Eleven of the twelve hybrids displayed a *K. ocellatus* mtDNA haplotype and only one a *K. hermaphroditus* haplotype, indicating that hybridisation is mostly between *K. ocellatus* hermaphrodites and the rare *K. hermaphroditus* males, given that no previous outcrossing has been observed between hermaphrodites (45, 46). The fact that five of the hybrids were backcrosses (four backcrosses with *K. ocellatus* and one with *K. hermaphroditus*), reveals that at least 50% of the hybrids are reproductively viable, despite high genomic divergence between the parental species. The asymmetry of these backcrosses could be influenced by the different mating systems of the parental species (predominant selfing vs outcrossing). The absence of F2 individuals may indicate that hybrids reproduce mostly via sexual backcrossing with a parental species, although it is unclear whether they can also mate between each other or self-fertilise.

Reproductive isolation increases with genetic distance between species pairs (47, 48) potentially due to postzygotic hybrid incompatibility caused by the gradual accumulation of divergent alleles. However, the hybridisation found here, which is comparable to the introgressive hybridisation between other highly divergent species (48, 49) indicates that even the offspring of very genetically divergent taxa can be viable. *Kryptolebias ocellatus* and *K. hermaphroditus* are very divergent genetically (50, 51) but also epigenetically. Although we cannot rule out that microecological differences between species (e. g. diet, habitat use) may have influenced their DNA methylation profiles, previous evidence of a close relationship between the genome and methylome in response to environmental variation (52, 53), suggest that the DNA methylation differences between *K. ocellatus* and *K. hermaphroditus* are primarily-driven by genomic differences between the species.

Most of the studies in plants using methylation-sensitive amplification fragment length (MS-AFLP) indicate prominent additive effects in cytosine methylation patterns, with typical Mendelian inheritance (35, 36, 54, 55) with few examples of non-additive effects in hybrids, potentially related to phenotypic plasticity and adaptation (56, 57). However, high-throughput sequencing has revealed the importance of non-additive DNA methylation states in crop hybrids (37, 38) which could alter gene expression levels (58). Thus, while most genes seem to be additively expressed in hybrids, there are still thousands of non-additive changes in transcript levels, some resulting from the modification of the epigenetic marks, which are likely involved in hybrid heterosis or incompatibility (59).

The strong intermediate effects observed in the methylation patterns between of *K. ocellatus* and *K. hermaphroditus* hybrids highlight the importance of the genetic background on DNA methylation levels. This evidence is reinforced by the DNA methylation patterns of the backcross individuals, more similar to the parental species than to the F1 hybrids. However, a small proportion of the DMCs between hybrids and the parental species displayed non-additive, over or underdominant effects in the hybrids, which could represent a mechanisms to generate phenotypic diversity (34), and potentially increased hybrid fitness (3). Although few studies have assessed DNA methylation inheritance in fish hybrids, the majority of DNA methylation effects appear to be additive, at least in allopolyploid hybrids (60, 61), as oberved here. This contrasts with transgressive patterns of gene expression found in some F1 fish hybrids (27, 30, 62), but generally agree with patterns found in hybrids among more genetically distant species such as *Drosophila* species and the house mouse, where divergent traits are regulated by more genes without a dominance pattern (31, 32).

The DMCs found between hybrids and both parental species may be expected to affect important biological processes many of them involved in developmental (i. e. central nervous system development, chordate embryonic development, eye development). As many of the hybrids are viable, these DMCs do not seem detrimental and could reflect allele-specific compensatory effects, with stabilising selection favouring an optimal level of gene expression by compensating the effects of single alleles through *cis* and *trans* regulatory factors (63). Compensatory effects in the gene expression of hybrids are well known (12, 27) but the existence of compensatory DNA methylation inheritance and epiallele dominance is not well understood (37).

Our study reveals that two genetically distant fish species with contrasting mating systems are capable of interbreeding and producing viable hybrids. Moreover, our results indicate that F1 hybrids are able to backcross with their parental species at high rates. Our results suggest that high hybridisation rates in this system might be favoured by natural selection because it creates additional genetic and epigenetic variation. Although the epigenetic effects were predominantly additive, we also found a small percentage of non-additive effects unrelated to the environment, which was common to both species. Such non-additive epigenetic variation could represent an evolutionary bet-hedging strategy to survive to environmental change, particularly in asexual or self-fertilising species.

## Material and methods

### Sample origin and microsatellite genotyping

In total, 103 *K. ocellatus* and 42 *K. hermaphroditus* individuals collected in locations from Sepetiba (Guaratiba, GUA) and Guanabara (Fundão, FUN) Bays, at the west and east limits, respectively, of the Rio de Janeiro municipality in Brazil (Figure S1) were genotyped for 16 microsatellites (39, 45). Sampling was carried out under license ICMBio/SISBIO 57145-1/2017 and approved by Swansea University Ethics Committee ref SU-Ethics-Student-250717/245. The final dataset consisted of a combination of previous datasets and new genotyping of fish colected in September 2017 using handnets (Table 1). Fish species were identified morphologically and confirmed by cytochrome oxidase subunit I (*cox1*) barcoding (39). Micro-checker v. 2.2 (van Oosterhout et al. 2004) was used to check for errors or presence of null alleles. FSTAT v. 2.9.3.2 (64) was used to measure departures from Hardy–Weinberg equilibrium.

STRUCTURE 2.3.4 (65) was used to identify genetic clusters with the following parameters: 10 iterations per K, a total of 1,000,000 MCMC and 100,000 burn-in, admixture model, independent allele frequencies, and testing K ranging 2–10. The most likely K was inferred using the likelihood ΔK method (66)) in Structure Harvester (67). Independent STRUCTURE runs were aligned and plotted using CLUMPAK (68). Genotypic associations among individuals were visualized using correspondence analysis (FCA) implemented in GENETIX v. 4.04 (69).

NEWHYBRIDS v. 1.1 (70) was used to estimate the posterior probability of each individual belonging to parental species, F1 hybrids, F2 or backcrosses between F1 and each parental species, based on their allele frequencies. The analysis was run using the default genotype proportions, uniform prior option, burn-in period of 50,000 iteration and 300,000 MCMC sweeps.

### Methylation-sensitive genotype-by sequencing library (msGBS)

To obtain genomic and cytosine methylation data simultaneously, we built a methylation-sensitive genotype-by sequencing library (msGBS) using pectoral-fin samples of 55 hermaphrodite individuals (33 *K.* ocellatus and 22 *K. hermaphroditus*) (Table S2). This protocol modifies the genotype-by-sequencing protocol described in Poland, Brown, Sorrells and Jannink (71) as shown in Kitimu, *et al.* (72). Genomic DNA was digested using a EcoRI and HpaII and ligated to barcoded adapters. A single library was produced by pooling 20ng of DNA from each sample and amplified in eight separate PCR reactions which were pooled after amplification, size-selected (range 200 – 350 bp) and sequenced in an Illumina NextSeq500 sequencer.

### Data processing

Paired-end reads were demultiplexed using GBSX v 1.3 (73). We then filtered (-qtrim r; -minlength 25) and merged the reads by individual using BBmap tools (74) mapped to *Kryptolebias marmoratus* reference genome (75) using Bowtie 2 v. 2.2.3 and generated filtered and indexed individual BAM files with samtools v. 1.9 (76).To call genotypes, we used ANGSD v 0.9.2.9 (77)). Single and double-tons were removed and, using SAMtools genotype likelihood model, we estimated posterior genotype probabilities assuming a uniform prior (-doPost 2). We also used the ANGSD (-SNP_pval 1e-6) to carry out a Likelihood Ratio Test to compare between the null (maf = 0) and alternative (estimated maf) hypotheses by using a X^2^ distribution with one degree of freedom. These analyses produced two genomic datasets for 53 samples: dataset I with 597,733 sites, coverage between 12.0X and 346.5X (mean 145.2X) and missing data ranging from 0% to 7.2% (mean 0.50%), and dataset II with 5,477 SNPs, coverage between 12.4X and 382.6X (mean 152.9X) and missing data ranging from 0% to 4.9% (mean 0.34%) (Table S2). A strong correlation (R^2^ = 0.93, p < 0.001) between the size of each the 3,073 scaffolds of the *K. marmoratus* reference genome and the number of SNPs from each scaffold indicated that the SNPs were evenly distributed throughout the reference genome.

### Population genetics and hybridisation analysis with SNPs data

We used ANGSD to compute the unfolded global estimate of the Site Frequency Spectrum (SFS) for dataset I to calculate the observed proportion of heterozygous sites (PHt) per sample (Korneliussen et al. 2013). The observed fraction of heterozygous sites was calculated as the ratio between the number of heterozygotes and the total number of sites with information. A pairwise genetic distance matrix was computed directly from the genotype likelihoods from dataset II (SNPs data) using ngsDist v.1.0.2 (78) and was then used for Multidimensional Scaling (MDS) using the R package *cmdscale*.

To estimate individual ancestries, we used ngsAdmix v. 3.2 (79) on dataset II, ranging K between 2-10 for 100 replicates using default parameters, except for tolerance for convergence (-tol 1e-6), log likelihood difference in 50 iterations (-tolLike50 1e-3), and a maximum number of EM iterations (-maxiter 10000).

To investigate the hybridisation between *K. ocellatus* and *K. hermaphroditus*, we used a subset of individuals from FUN and GUA for both species (39 individuals, 17 with *K. ocellatus cox1* haplotype, 22 with *K. hermaphroditus cox1* haplotype). We called SNPs with the same parameters described above using ANGSD with no missing data allowed, and selected those with the highest pairwise F_ST_ values between species. Pairs of SNPs with significant (LD) were removed and randomly replaced with other SNPs to complete a dataset of 200 SNPs (the upper limit of NEWHYBRIDS) with low-levels of LD and high FST values. We then run NEWHYBRIDS v.1.1 with the same parameters described for the microsatellites to investigate the posterior probability of each individual to belong to one of the six hybrid classes.

### Differentially methylated cytosines and hybrid cytosine methylation patterns

To investigate cytosine methylation patterns, we selected *K. ocellatus* and *K. hermaphroditus* individuals from the two mangroves with hybrids (GUA and FUN, Fig S1), resulting in 39 individuals (Table S2). Hybrids were classified into F1 or backcrosses with each one of the parental species based on SNP data. DMCs were identified using the R package msgbsR (Mayne et al. 2018). Individual restriction-digested reads were aligned to the reference genome and filtered out for correct cut sites and possible outliers. The function *diffMeth* was used to split data according to comparisons, normalise read counts according to library size and identify DMCs. We performed three comparisons: (1) *K. ocellatus* vs *K. hermaphroditus;* (2) hybrids vs *K. ocellatus;* and (3) hybrids vs *K. hermaphroditus*. Only loci with more than 1 count per million (CPM) reads in at least “n” individuals in each compared group, with “n” being determined by the group with the lowest number of samples in each comparison (11 in *K. ocellatus* vs *K. hermaphroditus;* seven in the comparisons including hybrids). DMCs were then filtered with a false discovery rate (FDR) of 0.01 and the logFC was retrieved to evaluate the intensity and direction of methylation changes. We generated a list of common DMCs (FDR <0.01) present in the comparisons between hybrids vs *K. ocellatus* and hybrids vs *K. hermaphroditus* and the normalised counts of these DMCs across all individuals was used for the downstream analysis.

To visualise overall variation in DMCs, we performed a multidimensional scale analysis (MDS) using Euclidean distance across all individuals. To compare DMCs profile across experimental groups using hierarchical clustering, normalised counts per DMC and individual were scaled and the differences in normalised counts for each site were estimated. Inheritance was considered potentially additive if normalised counts of DMCs between the parental species were intermediate in the hybrids and over or under dominant if normalised counts were higher or lower in hybrids compared to the parental species (Fig. S2).

### Genomic context and gene ontology enrichment analysis

Using the annotated *K. marmoratus* reference genome (Rhee et al. (2017), we identified the genomic context of the DMCs common to the two comparisons between hybrids and parental species, i.e. within gene body, promoter or intergenic region (Supplementary material). To identify differences in DNA methylation across different genomic contexts, we run MDS using DMCs from each group.

The annotated regions affected by these DMCs were used for the gene ontology enrichment analysis using zebrafish (*Danio rerio*) gene orthologs in PANTHER v. 11 (Mi et al. 2016). We searched for enrichment across biological process ontologies curated for zebrafish. Only genes which matched with the gene names annotated for zebrafish were included in the gene ontology analysis.

## Acknowledgements

We are grateful to Larissa Rodrigues, Benjamin Mayne and Kiflu Tesfamicael for help with bioinformatics pipelines. This work was supported by National Geographic/Waitt program [W461-16] and Conselho Nacional de Desenvolvimento Científico e Tecnológico (CNPq) [233161/2014-7].

## Data accessibility

FastaQC files for msGBS library can be accessed at NCBI (accession PRJNA563625). Sequence data will be deposited in GenBank and microsatellite data will be submitted to Dryad upon ms acceptance. All scripts used in the project are available at https://github.com/waldirmbf/BerbelFilho_etal_KryptolebiasHybridisation.

